# Pupil dilations prior to freely timed actions reflect the reported timing of conscious intention

**DOI:** 10.1101/2024.05.01.592070

**Authors:** Jake Gavenas, Aaron Schurger, Uri Maoz

## Abstract

Freely timed actions are typically preceded by a slow anticipatory buildup of cortical brain activity, which has been extensively studied. However, such free actions are also preceded by slow pupil dilations in both humans and other animals, which have barely been examined. We investigated the neurocognitive significance of antecedent pupil dilations (APDs) in a voluntary-action paradigm. Participants performed self-paced actions and reported the timing of movement, conscious intention, or other events using a clock. APDs began a second or more before movement, and control conditions suggest that they did not reflect processing related to reporting demands, motor execution, or general anticipation. Critically, APD timing covaried with the reported timing of intention awareness but did not covary with the reported timing of overt movement or an external stimulus. Furthermore, decoding algorithms could distinguish APDs with above-chance accuracy more than 500 milliseconds before button-press. Our results suggest that APDs reflect a shift in awareness prior to movement onset and potentially offer a non-invasive method of predicting spontaneous movements before they occur.

Highlights:

1. Freely timed movements are preceded by antecedent pupil dilations (APDs).

2. APDs do not reflect reporting, motor execution, or general anticipation.

3. APDs are informative of upcoming movements 500+ milliseconds before button-press.

4. APD timing specifically correlates with timing of intention awareness.

## Introduction

The capacity to initiate actions spontaneously is fundamental to adaptive goal-directed behavior. Human and animal neuroscience has begun elucidating the neuronal substrates of voluntary action by investigating precursors of freely-timed actions^1–7^. Studies in humans have found that spontaneous voluntary actions are preceded by gradual buildups of neuronal activity in frontal regions such as the (pre-)supplementary motor area (SMA), anterior cingulate cortex, and motor cortex^4,8–11^. Similar anticipatory buildup signals in analogous regions have been reported in other animals prior to spontaneous or self-paced movements^6,7,12–14^.

A great deal of research has been devoted to elucidating the cognitive significance of these signals. Notably, similar anticipatory buildups have been observed in signals reflecting subcortical arousal mechanisms. In particular, several studies have found that freely-timed movements are preceded by pupil dilations in humans^15^ and other animals^16–18^. However, the neurocognitive significance of these antecedent pupil dilations remains poorly understood.

Pupil dilations have been linked to a variety of cognitive processes, including attention, cognitive effort, perception, decision-making, awareness, and memory encoding and recall^19–28^. Widespread reports of associations between pupil dilations and cognitive processing likely stems from the well-documented relationship between pupil size and subcortical neuromodulatory hubs, such as the locus coeruleus^16,29–31^, which are themselves likely involved in myriad cognitive functions. Crucially, pupil dilations are particularly sensitive to changes in awareness^19,23,32–34^. Furthermore, gradual pupil dilations like those observed before spontaneous movements are also observed before the generation of creative ideas^35^, eureka moments during problem solving^36^, free recall^28^, and switches during bistable perception^37,38^, which suggests they may reflect processing related to shifts in awareness. In particular, Salvi and colleagues recently suggested^36^ that gradual pupil dilations before eureka moments during problem solving reflect the “switch into awareness” of a solution (or the restructuring of information into a conscious thought), which may have some commonalities with spontaneous voluntary action in terms of its underlying neural mechanisms^39^. If that is the case, then pupil dilation timing should specifically covary with the timing of subjective experience. More specifically for spontaneous voluntary action, we speculated that pupil dilations might covary with the subjective experience of intention onset, more so than with other peri-movement-onset phenomena.

We hypothesized that antecedent pupil dilations (APDs) specifically relate to the conscious decision or intention to initiate movement *before* the onset of voluntary action. To test this hypothesis, we recorded pupil size from human participants during a voluntary-action paradigm, in which participants reported the timing of their subjective urge or intention to move using a clock^2^. Specifically, participants moved at a time of their choice; then they either did not report anything, reported the timing of their movement, or reported the timing of their decision to move. On other trials, participants imagined moving and reported the timing of their imagined movement or listened for an auditory stimulus and reported its timing without initiating action. We aimed to answer three questions in this study: (1) Do APDs specifically reflect spontaneous shifts in awareness, rather than other cognitive processes, such as allocating attention for later reporting, motor execution, or general anticipation? (2) When do APDs begin and are they predictive of upcoming movements? (3) Does the timing of APDs reflect the reported timing of conscious intention? We found that the presence of dilations was unrelated to motor execution, reporting demands, and general expectation. Furthermore, decoding analyses suggest that APDs are distinguishable with above-chance accuracy 500+ milliseconds prior to action, when time-locked to (and hence conditioned on) action onset. Finally, the timing of dilations was related to the reported timing of the urge or intention to move but not to the reported timing of the movement itself or to that of an external stimulus. Our results therefore provide evidence that antecedent pupil dilations indeed reflect the “switch” into awareness of a subjective decision to move prior to movement initiation.

## Results

Participants (N=29) completed a voluntary action task while reporting their internal state using a clock. The procedure is described in Figure 1, and details are given in Methods. Participants were instructed to wait for the clock to make one full revolution (2.5 sec) and, after that, press the spacebar whenever they felt like it. They were further instructed to press spontaneously and not pre-plan their actions. In the no-report condition, participants just pressed the space bar and did not report anything. In the other conditions, they reported when they moved (M-Time), when they felt the urge or intention to move (W-Time), when they imagined moving (I-Time), or when they heard the randomly occurring tone (S-Time), using the clock. They reported this by clicking the location on the clock corresponding to the onset of the event.

**Figure 1:**
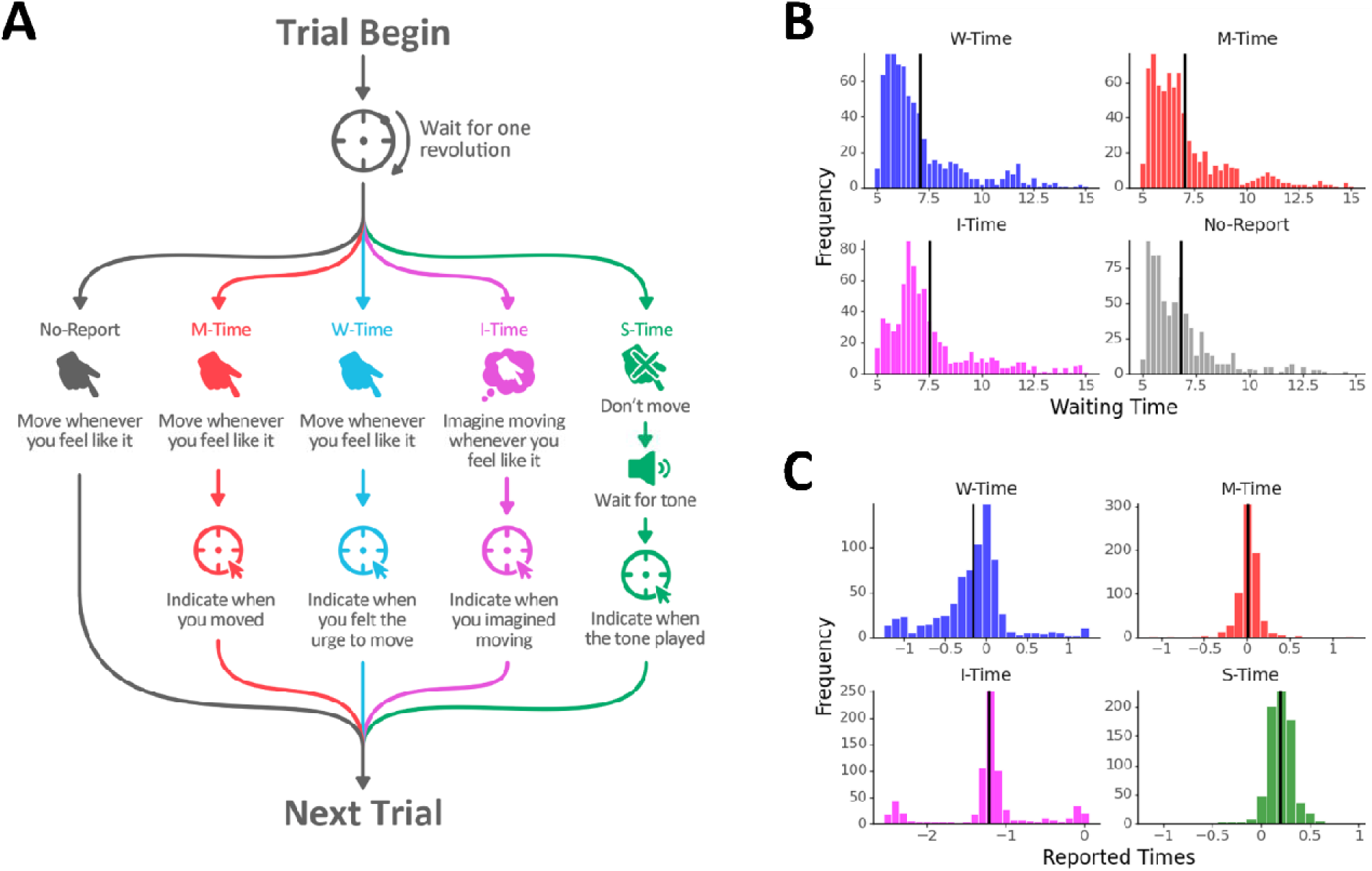
Paradigm overview & behavioral results. Participants completed a self-paced action task with 5 different conditions, organized in a blocked manner (see Procedure in Methods). Participants initiated the trial at a time of their choosing by pressing the spacebar when they were ready. At trial onset, a clock would appear onscreen with a dot located at the top of the clock. The dot began rotating at a rate of 1 cycle per 2.5 seconds. Participants were instructed to wait for one cycle and then either press the spacebar whenever they felt like it (No-Report, M-Time, W-Time conditions); imagine pressing the spacebar whenever they felt like it, noting the time on the clock, and then pressing the spacebar after an additional half revolution had elapsed (I-Time condition); or to avoid moving and just wait for a brief tone to play (S-Time condition) and note the time of the tone on the clock. After they had pressed the spacebar or the tone had played, the dot on the clock would finish its current revolution, make one more revolution and then disappear. The participants would then either move on to the next trial (No-Report) or make their report according to task demands—W-Time (report the dot’s location at the onset of their urge or intention to move), M-Time (report the dot’s location at the onset of their actual movement), I-Time (report the dot’s location at the onset of imagining the movement), or S-Time (report the dot’s location at the onset of the tone). After this they would continue to the next trial. **B:** Histogram (pooled across participants) of waiting times until button press relative to trial start for the W, M, I, and No-Report conditions. Black vertical lines are condition-specific averages. **C:** Histogram (pooled across participants) of reported timings of the urge or intention to move (W), movement (M), imagined movement (I— note that we only retained trials with I reports between-2.0 and-0.5 s for further analysis), or stimulus (S) relative to button press or tone onset. Black vertical lines are again condition-specific averages.

The W-Time condition was the main target of the experiment. The M-Time condition served as a control for externally directed awareness of action. The S-Time condition controlled for general anticipation or expectation effects. Participants did not have to move following tone onset in the S-Time condition. The I-Time condition controlled for any effects of motor execution (as participants imagine moving spontaneously and then pressed the spacebar after half a clock revolution to end the trial). And the No-Report condition controlled for any effects of allocating attention in order to make their report.

Participants completed practice blocks of the No-Report condition first, then completed half of the No-Report trials. After that they practiced the other four conditions (W-Time, M-Time, I-Time, S-Time), where reporting was required, in randomized order. Next, they completed multiple blocks of the four reporting conditions, in randomized order in a blocked design. Finally, they completed the second half of the No-Report trials.

### Behavior

Participants waited around 7 seconds to move or imagine moving on average (Fig. 1B; means & 95% confidence intervals across participants: **No-Repor**t: 7.024 s, [6.462, 7.587]. **W-Time**: 7.310 s, [6.747, 7.872]. **M-Time**: 7.207 s, [6.645, 7.770]. **I-Time**: 7.857 s, [7.294, 8.419]; obtained via Linear Mixed-Effects (LME) analysis). Waiting time in the I-Time condition was significantly longer than the other conditions, perhaps due to the added task demands of imagining a movement (post-hoc tests from LME analysis; I vs. no-report: t(2943.942) = 6.703; p < 0.001. I vs. W: t(2943.942) = 4.404; p < 0.001. I vs. M: t(2943.942) = 5.228; p < 0.001). Waiting times in the No-Report, W-Time, and M-Time conditions were not significantly different.

Participants’ timing reports were in line with prior results, with W-Time being roughly 150 milliseconds before movement, M roughly at the time of movement, and S approximately 200 milliseconds after tone onset (Fig. 1C; participant-specific means & 95% confidence intervals across participants: **W-Time**:-0.155 s, [-0.185,-0.125]. **M-Time**: 0.014 s, [-0.016, 0.044]. **S-Time**: 0.194 s, [0.164, 0.224]; obtained via Linear Mixed-Effects). W and S reports were significantly earlier and later than zero, respectively (both p < 0.001), whereas M reports were not significantly different from zero (p = 0.348). Differences between reported W, M, and S-times were all highly significant (all p < 0.001). In the I-time condition, participants reported that they imagined moving 1.217 seconds [1.183, 1.251] before recorded button presses, consistent with a task demand to press ½ a clock rotation (i.e., 1.25 seconds) after spontaneously imagining moving. In some trials participants reported imagining moving close to movement onset or a full clock rotation before movement (see Fig. 1C), suggesting lapses in attention. We retained only trials with I-Time reports between-2 and-0.5 s for further analyses.

### Antecedent pupil dilations (APDs)

Spontaneous movements were preceded by gradual APDs beginning 0.5-1 s before movement in the No-report, W-Time, M-Time, and I-Time conditions, whereas passively experienced yet generally anticipated tones (S-Time condition) were not preceded by such dilations (Fig. 2, individual participants’ dilations in Fig. S1). Their absence from the S-Time condition suggests they do not reflect general anticipation, because participants knew a sound was going to occur. In contrast, their presence during No-Report trials suggests they are not specifically tied to allocating attention for reporting. Furthermore, their presence during imagined movement suggests they do not reflect processes related specifically to motor execution.

**Figure 2:**
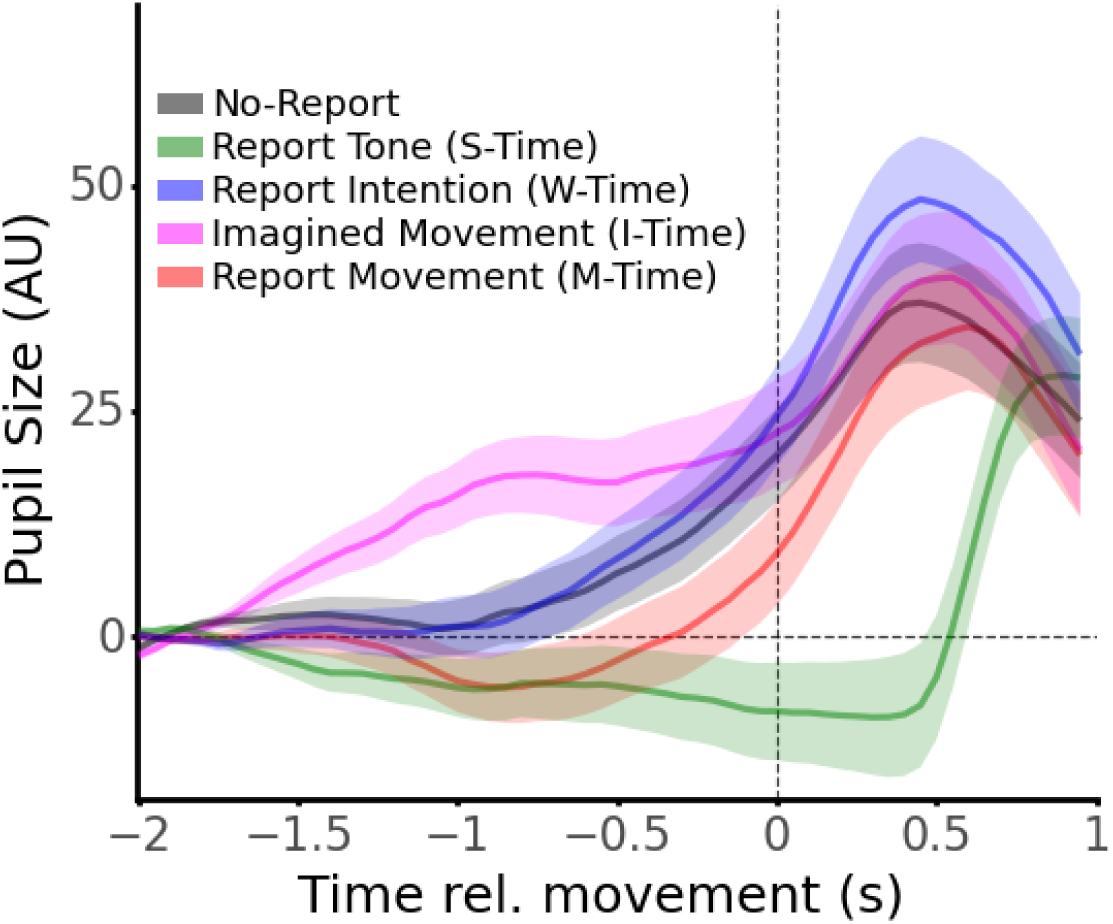
Pupil dilations before spontaneous actions. Pupil size (arbitrary units, AU) relative to the time of button-press (No-Report, W-Time, M-Time), imagined movement (I-Time), or tone onset (S-Time). Solid lines are average pupil size for each condition and shaded regions are standard error (both obtained by fitting an LME model to data at each time point; see Methods for analysis details & preprocessing steps). For the No-Report, W-Time, and M-Time conditions, t=0 signifies the time of recorded button press. For the S-Time condition, t=0 signifies the time of tone onset. For the I-time condition, t=0 signifies the time of imagined movement, reported post-hoc using the clock. Pupil dilations preceded spontaneous movements and imagined movements but did not precede tones. Dilations did not much depend on the need to report (No report vs. W-Time & M-Time conditions) and did not require motor execution at t=0 because they also occurred before imagined movements (I-Time condition).

Furthermore, APDs were present prior to imagined movements (I-Time condition), where we aligned the data to the reported times of imagined movement (see Methods). The pupil waveform in the I-Time condition reached a maximum dilation size that are visually of similar magnitude to the conditions in which participants made overt movements, which again suggests that the dilations do not reflect motor execution. Notably, dilations before imagined movements had a somewhat different early waveform than conditions with overt movement. This is possibly due to the relative uncertainty in the exact timing of the imagined movement (in comparison to overt movements).

In the S-Time condition, the participants waited for the tone without moving. So, this condition involved no spontaneous movement. As expected, the pupil waveforms in this condition were therefore markedly different than in the other conditions, remaining roughly flat until close to 500 ms after tone onset, where the pupil dilated rapidly to a size more similar to its peak size in the other condition. Hence, pupil waveforms had a significantly greater slope (i.e., stronger dilations) in W-Time, M-Time, I-Time, and No-Report conditions compared to S conditions in the 1.5 seconds before movement onset (p_Tukey_ < 0.001 for W-Time, I-Time, and No-Report versus S-Time, p_Tukey_ = 0.008 for M-Time versus S-Time; Linear Mixed-Effects). These results suggest that APDs do not reflect processing related to motor execution or general anticipation.

### APDs are informative of upcoming movements

To investigate the timing of APD onset, we employed a breakpoint analysis using a model-comparison approach. Briefly, we fit APDs (obtained from trials with overt spontaneous movements, i.e. No-Report, W-Time, and M-Time trials, so that ground truth of movement onset was known) with models where a flat trend (or fixed value) would continue until a “breakpoint,” after which the model could increase linearly, quadratically, or exponentially. We fit the models with breakpoint times between-1.5 seconds to +0.15 seconds relative to movement onset (fitted on data between-2 seconds and +0.2 seconds; baselined using mean pupil size in the range [-2,-1.5] for reach trial; fitting on non-baselined data resulted in largely the same results). We then extracted the Akaike Information Criterion (AIC), a quantifier of a model’s goodness-of-fit (see Breakpoint Analysis in Methods for Details). We found that the best performing model was the quadratic model with a breakpoint at 1.0 seconds before movement onset (Fig. 3A; best performing models: Linear: AIC= 530134.925 at-0.65 s; Quadratic: AIC= 530110.624 at-1.0 s; Exponential: AIC= 530116.876 at-0.8 s). This model was a good fit for the average pupil dilation prior to movement (Fig. 3B). Relaxing the requirement that the models must have a constant value before the breakpoint, and allowing a linear trend instead, resulted in a better fit (lower AIC), with slightly later estimated dilation onsets (Fig. S2A). The best performing model among these three was the linear-then-quadratic model, with an onset of 0.9 seconds before movement onset (best performing models: linear-then-linear: AIC= 530123.903 at-0.55 s; linear-then-quadratic: AIC= 530107.562 at-0.9 s; linear-then-exponential: AIC= 530113.437 at-0.7 s). However, these models did a poorer job of visually capturing a specific moment of dilation onset (see Fig. S2B), and the differences between AIC for the best performing models of each class was very small (3 out of ∼530,000), so it is not clear that the minute decrease in AIC is worth the more complex model.

**Figure 3:**
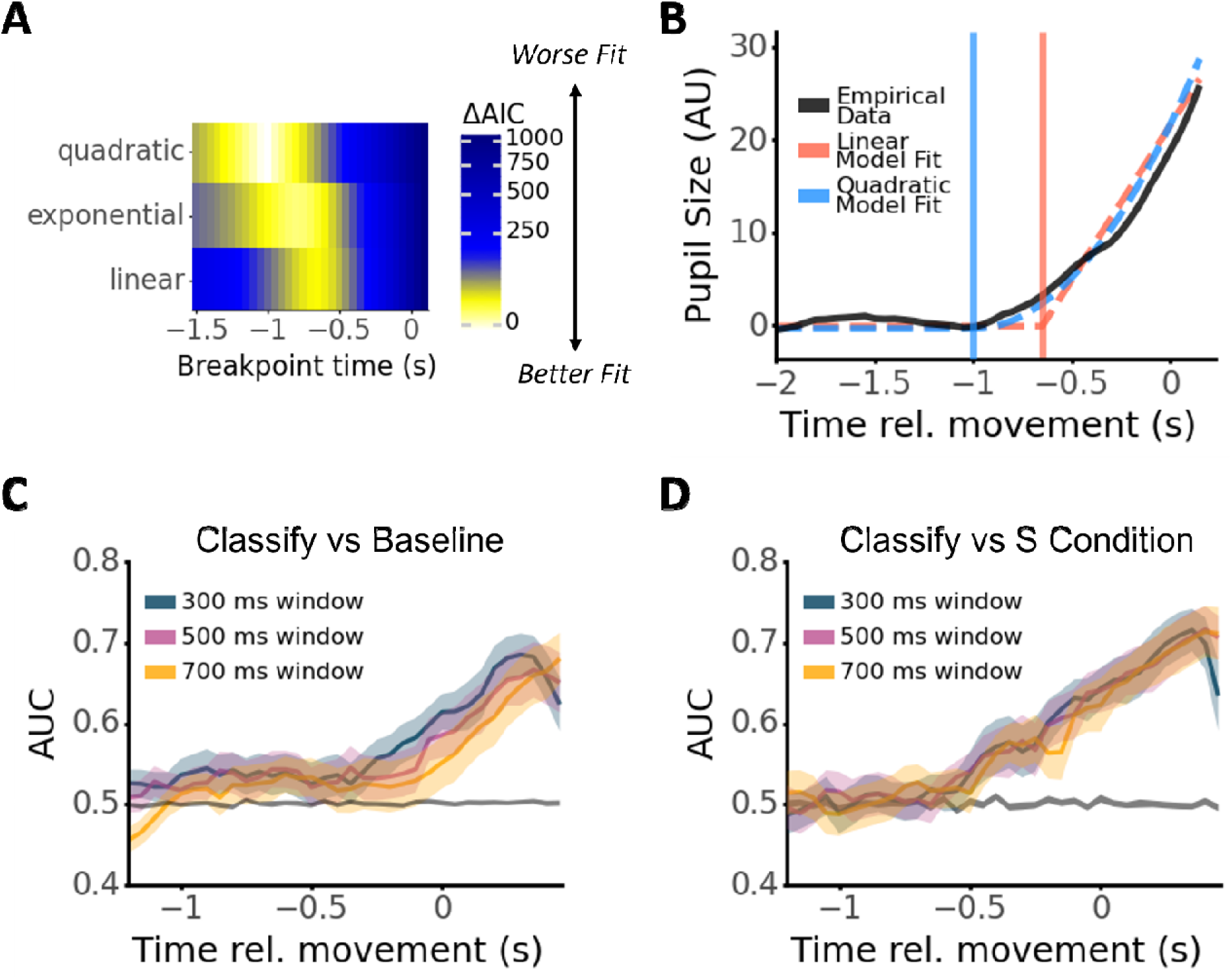
Characterizing the onset of APDs. A: Breakpoint analysis. We fit several piecewise models to the APD data, which included a constant value up to the breakpoint and then either a quadratic, exponential, or linear increase following the breakpoint (for simplicity, piecewise models are referred to as quadratic, exponential, and linear). The breakpoint range was between-1.5s and 0.2s relative to movement onset. The colors designate the difference in AIC values from the AIC of the best-performing model (lower AIC is associated with a better model fit). The best performing model overall was a quadratic model with breakpoint at-1.3s. **B:** Model fits using the breakpoint that resulted in lowest AIC for the linear and quadratic models (exponential omitted due to overlap with quadratic model). Solid black line is grand-average pupil size (averaged across trials & W, M, and No-Report conditions, and then across participants), dashed lines are model fits, and vertical solid lines indicate the breakpoint used for the corresponding model. **C-D:** Decoding analysis. Test-set AUC (area under the ROC curve; measure of machine-learning classifier performance; average of 3-fold cross-validation) when classifying pupil slope in the spontaneous movement conditions (W, M, and No-Report) at each time point from pupil slope during a baseline period (**C**) or at an equal time relative to tone onset in the S-Time condition (**D**). Slopes were calculated in sliding window (varying size; 50 ms step), where time on the x-axis refers to each window’s leading edge (latest time-point). Solid lines are mean AUC across participants, shaded regions are standard error across participants. Dark shaded regions around AUC = 0.5 reflect chance-level AUCs (standard error above and below mean) obtained from 100 shuffles of the data pooled across participants (to allow calculation of AUC in all shuffles). Decoding AUC began rising above chance around ∼0.5 seconds and ∼0.7 s before movement onset when decoding versus a baseline and versus the S condition, respectively.

The breakpoint analysis suggests APDs begin relatively early before movement. However, it has been demonstrated that aligning and averaging autocorrelated signals to movement onset may introduce artifacts, including slow ramping signals towards the onset of the movement (also because in such cases movement onset is statistically dependent on the neural activity preceding it)^14,40–42^. It is therefore not clear whether the dilations in Fig. 2 are *predictive* of an upcoming spontaneous movement. To investigate this, we used a decoding approach: we trained machine-learning classifiers (linear discriminant analysis—see Decoding Analysis in Methods) to discriminate pupil slope in a sliding window from slope during a baseline period (Fig. 3C) (leading edges in Fig. 3C-D; performance on individual participants in Fig. S3). Here we focused on conditions with overt spontaneous movements (No-Report, W-Time, and M-Time) from slopes during a baseline period (−2 seconds to-2 + window size). We analyzed pupil slope rather than pupil size to avoid introducing potential confounds due to our choice of baseline, due to differences in tonic pupil size across trials, or due to slow drift in the pupil signal. Decoding accuracy hovered around chance until ∼500 milliseconds before movement, after which it started rising, reaching a test-set AUC of 0.619 at movement onset (window size 0.3 s; 3-fold cross-validation; Fig. 3C), after which decoding performance kept increasing, reaching a maximum AUC of 0.685 at 0.3 seconds after movement onset, and then dropped off (presumably because the pupil begins constricting following dilation—see Fig. 2). We also trained another LDA classifier to discriminate pupil slopes in the conditions with spontaneous movements (No-Report, W, and M) from the pupil slope in the S-Time condition (which included no movement, just passive listening and attending)—at matched time points relative to movement/tone onset (see Decoding Analysis in Methods). This method avoids introducing confounds due to baseline correction (see refs.^42,43^). Decoding performance was now at or near chance until ∼700 milliseconds before movement, when it started rising, reaching a test-set AUC of 0.645 at movement onset (window size 0.3 s; 3-fold cross-validation; Fig. 3D) and a maximum AUC of 0.715 at 0.35 seconds after movement onset. These increases in AUC were accompanied by clear shifts in the distribution of pupil slopes in the positive direction at times closer to movement, indicating pupil dilation (Fig. S4). Taken together, these analyses suggest that APDs show a non-stationarity that is not due to baseline correction between 500 and 700 milliseconds before movement.

### Timing of APDs specifically relates to the reported timing of intention awareness

We next investigated the cognitive significance of APD timing. Specifically, we tested the hypothesis that the timing of APDs was related to the reported timing of awareness for the W-Time, M-Time, and S-time conditions. Following prior studies of how movement-preceding signals relate to participants’ reports^44–46^, we performed a median split of the pupil data according to each participants’ reported W-times. We found that earlier W-times were accompanied by earlier and stronger dilations (Fig. 4A). A cluster permutation test suggested the correlation between dilations and W-times was significant (p = 0.036; see Analyzing Dilations & Subjective Reports in Methods). We further tested this finding by repeating the analysis with more strict exclusionary criteria and still found that earlier W-times were associated with earlier dilations (omitting participants with fewer than 10 W-trials; N=14; p < 0.01 non-parametric cluster permutation test, 100 bootstraps; Fig. S5A). Recreating this analysis on non-baselined data did not result in significant differences, due to variance in pupil size across trials. But it did show that dilations on early W trials reach a higher peak pupil size compared to late W trials (Fig. S5B). Demeaning the pupil data by subtracting whole-trial averages also suggested earlier and stronger dilations for earlier W-times (Fig. S5C). We verified this difference in dilation timing by repeating the breakpoint analysis described in the prior section separately for trials corresponding to “early” versus “late” W-times (within-participant median split), and confirmed that earlier W-times were associated with earlier dilations (best-fitting model was Exponential with breakpoint at-1.75 s; AIC = 89686.313; Fig. 4B) compared to later W-times (best-fitting model was Quadratic with breakpoint at-0.85 s; AIC = 82895.976). These results suggest a reliable relation between the onset of pupil dilations in spontaneous action and the W-times that participants reported.

**Figure 4:**
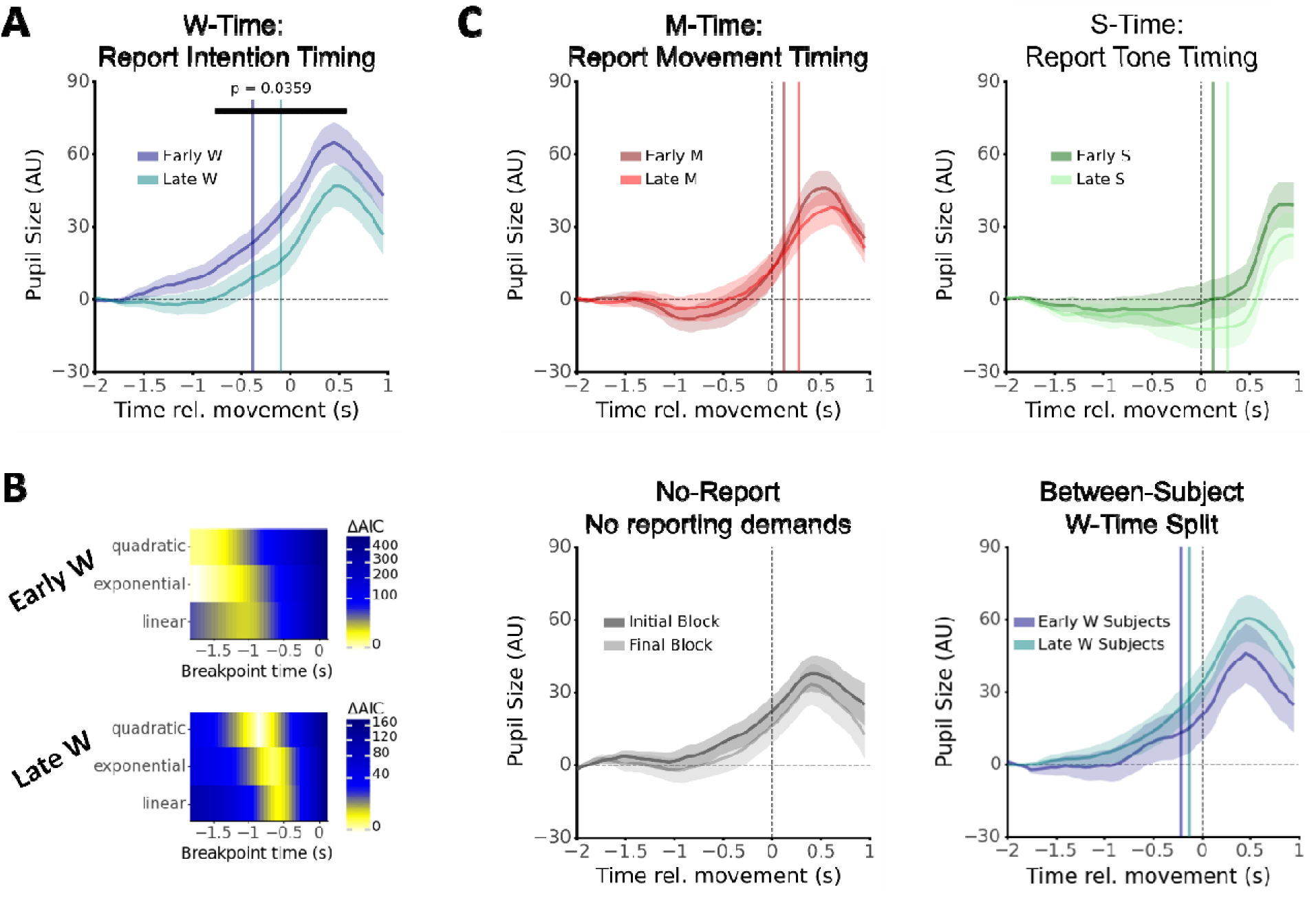
Timing of antecedent pupil dilations reflect self-reported timing of conscious intention onset. A. Mean & standard error of pupil trajectory over time (estimated using LME analysis) for relatively early and late W-times (within-participant median split). Colored vertical lines are average “early” and “late” W-times. Horizontal black line shows time of significant difference between trajectories (p < 0.05 from LME analysis). Shown p-value was obtained from distribution of largest-continuous-cluster obtained from shuffled data (N=1000 bootstraps). **B.** Breakpoint analysis on pupil trajectories for “early” and “late” W-time trials separately (otherwise as in Fig. 3). **C.** Means & standard error of pupil trajectory when splitting data according to reports of movement onsets (M-Time, within-participants median split, upper left), tone onsets (S-Time, within-participants median split, upper right), whether the dilations were in the initial or Final No-Report block (within-participants split, bottom left), or whether participants were “early” or “late” W-time reporters (W-time, between-participants split, bottom right). In all of these cases we did not find significant differences in pupil sizes. Vertical lines correspond to mean timings for early and late reports (upper left, right, and bottom right).

To test whether the relation between APD timing and subjective reports was specific to the W-time condition, we conducted the same median-split analysis as above on M-Time and S-Time conditions. We found no relation between M and S times and dilation timing (Fig. 4C; there were no significant clusters prior to movement/tone onset when splitting according to M or S times, so p-values could not be calculated). Furthermore, we investigated the effects of learning and fatigue on APDs by comparing APDs during the first block of No-Report trials (when participants did not yet know about all the other conditions—see Procedure in the Methods) to the last block of No-Report trials (at the end of the experiment). We found no reliable difference in APDs when comparing the two blocks (Fig 4C bottom left). Finally, we investigated whether participants who reported earlier W-Times also had earlier pupil dilations by performing a between-participants median split on W-times and comparing pupil trajectories as above. However, we found no such relationship (Fig. 4C bottom right), suggesting that this is primarily a within-participants effect.

## Discussion

We set out to investigate three questions: (1) Do antecedent pupil dilations (APDs) specifically reflect spontaneous shifts in awareness, rather than other cognitive processes, such as allocating attention for later reporting, motor execution, or general anticipation? (2) When do APDs begin and are they predictive of upcoming movements? (3) Does the timing of APDs reflect the reported timing of conscious intention? We recorded pupil size from participants while they made freely-timed voluntary movements and then reported the timing of their awareness of various events using a clock^2^. We found APDs before actual and imagined movements, but not before anticipated auditory stimuli. The presence of APDs on No-report and on imagined movements (I-Time) suggest that they do not reflect processing related to reporting or motor execution, respectively. The absence of APDs when reporting the onset of a tone (S-Time) suggests that they do not reflect processing related to general anticipation. Hence, in relation to our first research question, we concluded that APDs specifically reflect spontaneous shifts in awareness. For our second research question, we found that machine-learning decoding algorithms could classify whether APDs were occurring with above-chance accuracy 500-700 milliseconds before movement onset, compared to baseline pupil slope or to time-matched pupil slope before an auditory stimulus. This suggests that the early segment of APDs are not a result of backward-averaging the autocorrelated pupil waveform, time-locked to movement onset as has been claimed in the case of other pre-movement signals, such as the readiness potential^14^ (RP). Finally, in relation to our third research question, we found that earlier APDs were significantly associated with reported times of the intention to move (W-Time) but not with reported times of movement (M-Time) or stimulus (S-Time), suggesting that pre-movement dilations are specifically related to the onset of the intention to move.

Our study builds on a large body of work investigating the physiological precursors of voluntary actions. While most prior studies in this literature focus on cortical precursors of movement, we investigated pupil dilations, which presumably reflect activity in subcortical regions. In particular, changes in pupil size under constant luminance are closely related to activity in the Locus Coeruleus^29,30^ (LC). The LC is a subcortical neuromodulatory hub that releases norepinephrine to cortical and subcortical targets in response to surprising, conflicting, or other types of stimuli^47^. Norepinephrine release underlies shifts in attention^19,48–50^ and is related to changes in awareness-related states, such as from sleep to waking^51^. Norepinephrine inhibitors also decrease the frequency of spontaneous locomotion in mice^52^, suggesting that LC activity and norepinephrine release may facilitate action initiation.

However, the mechanism through which LC activity relates to cortical precursors of voluntary movements, such as the RP, remains unclear. In this respect, it is worth noting that the RP is thought to originate from the supplementary motor area (SMA)^10,14^, a sub-region of the medial frontal cortex (MFC). MFC also shows slow-ramping activity prior to voluntary action in fMRI BOLD signal and in single-neuron recordings^4,8–10^. Prior work has proposed that spontaneous voluntary movements are triggered when a weak drift-diffusion process in MFC crosses a threshold, and that the RP and other slow-ramping signals reflect that diffusion process when aligned to threshold-crossing^40–43,53^. Notably, MFC has strong reciprocal connectivity with the LC^54,55^, and is likely the only cortical region that projects to LC^56^. One possible mechanism tying our results to prior models is therefore that threshold-crossings by MFC activity fluctuations trigger LC activation and norepinephrine release due to recurrent excitation between these two regions, leading to both the ‘switch into awareness’ (which has previously been linked to threshold-crossings^57^) of the intention to move as well as facilitating motor execution^41^. Note that this proposed mechanism suggests that threshold-crossing occurs around or soon after the time when APDs become distinguishable via machine-learning methods, ∼500 ms prior to movement onset. While plausible, further research is needed to investigate this hypothesis.

We also found that machine-learning algorithms could distinguish APDs on no-report, M, and W trials from baseline on those trials and time-matched pupil waveform on S trials with above-chance accuracy beginning between 700-500 ms prior to button-press. Although our decoding performance at that time was only slightly above chance, the timing of increasing accuracy is comparable with studies using non-invasive EEG to predict upcoming movement. For instance, Bai and colleagues could predict an upcoming movement on average 620 ms prior to movement onset (with accuracies ranging between 57% and 90% depending on the participant)^58^. A more optimized decoding pipeline could potentially increase the accuracy of decoding based on pupil dilations (e.g. by using both eyes simultaneously). Given that acquiring decent-quality pupil size is simpler than acquiring decent-quality EEG data, this opens new possibilities for real-time prediction of voluntary movement initiation. Interestingly, Lew and colleagues^59^ were able to distinguish pre-movement activity from a baseline above-chance more than 1 second prior to movement by recording intracranially from contra-and ipsilateral SMA. But their accuracy remained roughly flat at a value only slightly above chance, until around 1 second before movement, after which it began increasing. These findings and ours suggest a specific event occurs 500-700 ms before movement that leads to a non-stationarity in their EEG data and our pupil data that drives increasing decoding performance leading up to movement.

Our results also bear on investigations into the timing of conscious intention (W-time) in relation to voluntary movements and neural precursors of action^2,60,61^. The validity of W-time reports obtained using the clock method as a measure of intention onset has been questioned due to several findings. First, W-time supposedly reflects an event (decision onset) that fully takes place before movement onset, but W-time reports were shown to be biased by events that occur after the movement^62,63^. Second, several studies have investigated potential relations between the timing of neural precursors of voluntary action, such as the RP, and W-Time. However, to our knowledge, none have ever been found^44–46,64^. Third, W-time seems to suffer from order effects and may be reported before movement solely due to task demands^65^. Based on these results, some have cautioned against using W-time as an index of the awareness of decision or will to move, with some suggesting that W-time reports may be retrospectively inferred based on movement timing and other factors rather than directly perceived prior to movement^60,63,65,66^. However, our finding that W-time is correlated with APDs, a pre-movement signal, suggests that W-time reports may not be entirely retrospective. Instead, W-time may emerge from an integration of prospective and retrospective factors^60,67^. Importantly, our results also suggest that APDs could offer a covert, non-invasive method for timing conscious intentions. This method could be of use in healthy populations, alongside more traditional reporting methods, but also in other human populations that cannot readily report, such as infants and locked-in patients, as well as non-human animals.

Although we investigated pupil dilations before spontaneous voluntary actions, pupil dilations are also observed before other types of spontaneous free behavior, including eureka moments during problem-solving^36^, creative idea generation^35^, free recall of memories^28^, and conscious switches during perceptual bistability^37,38^. In particular, Salvi and colleagues^36^ suggested that pupil dilations before eureka moments reflected the “switch into awareness” of a solution via reorganization of information into a new conscious percept. Our results favor their suggestion, especially considering that different types of spontaneous mental events are hypothesized to occur via a common mechanism^39,68,69^. That mechanism may itself be related to the circuitry that elicits APDs. The hypothesis that APDs reflect the “switch into awareness” across different types of spontaneous behaviors is further evidenced by findings that conscious decisions, but not the actions that express those decisions, are accompanied by pupil dilations^27^. Notably, similar slow-ramping buildups of neural activity precede other types of spontaneous behavior, including creative idea generation^70^, eureka moments during problem solving^71^, and free recall^69,72^, which may reflect spontaneous fluctuations that trigger a thought or action upon crossing a threshold^39,42^. The similarities between volition and other spontaneous behavior are striking and, we think, deserve further exploration.

Notably, our study was limited in a few ways that future studies might improve on. Foremost, we only recorded pupil size and were therefore unable to assess whether APDs are related to other signals—such as the RP—directly. Future studies may remedy this via simultaneous EEG and pupillometry recordings. Furthermore, the circuitry underlying shifts in pupil size is complex and pupil size is likely an imperfect index of LC activity^16,30,73^. Future studies could resolve this issue by recording pupil size as well as intracranially from the LC (for example in an animal model). Finally, although our findings in the S-Time condition provide evidence that APDs do not reflect general anticipation, our results do not rule out the possibility that they reflect anticipation of an event at a particular time. However, such specific anticipation may itself be related to the timing of conscious intention in the case of spontaneous voluntary action.

Taken together, our results have important implications for theorizing about conscious volition, for the interpretation of prior results relating to slow ramping signals (such as the RP) and how they relate to prospective awareness of motor intention, and for the possibility that antecedent dilations may reflect the switch into awareness for spontaneous thoughts in other contexts. Future studies might investigate whether the timing of APDs is also associated with the timing of subjective experience in the context of other spontaneous mental events, such as free recall and problem-solving via insight.

## Methods

### Participants

We recruited 37 participants from the Chapman University undergraduate population to participate in our study (mean age: 19.09±1.33 (stdev) for 33 participants; the age of 4 participants, who took part prior to the long university closures due to COVID-19 pandemic, was unrecoverable; 8 identified male, 29 female). Eight participants were excluded due to technical issues (mainly poor performance and data omissions by the eye-tracker), so our study encompasses results obtained from 29 individuals.

### Procedure

Prior to the experiment, participants provided informed consent (all study procedures were approved by the Chapman University ethics committee, IRB-20-122). Participants then sat at a desk 85 cm from the computer screen, under dim light conditions. After calibrating the eye-tracker, the participants were given instructions for the No-Report condition (see below). They then practiced the task for 10 trials. After that, the participants completed half of the No-Report trials (10 for participants 1-12, 15 for participants 13-29). They then completed training blocks for the W-Time, M-Time, S-Time, and I-Time conditions (10 trials each), with experimenters giving instructions for each condition when the corresponding training block began. Training blocks were presented in random order. After training had completed, participants completed 2 (participants 1-12) or 3 (participants 13-29) blocks of 10 trials for each of the conditions. Abbreviated instructions were provided at the beginning of each trial to ensure that participants were adhering to task demands (verified by behavior, Fig. S1), and blocks were delivered in randomized order. After completing this section of the main experiment, participants completed a final block of No-Report trials (splitting the No-Report trials in this way allowed us to see if there was an effect of training on the APDs—and there wasn’t, see Fig. 3D).

For all conditions, the participants were instructed to fixate on a dot in the center of a clock that was 5 cm in diameter (hence at ∼3.37 degrees visual angle; clock and fixation dot were white on a gray screen). A small white dot was shown revolving around the clock at a rate of 2.5 seconds per revolution (revolution speed in line with prior experiments, e.g., Dominik et al., 2018, Fig. 1). Participants were instructed to maintain fixation on the small fixation dot at the center of the clock while paying attention to the location of the other, rotating dot. The clock was designed to be small enough on-screen to make it easy to keep track of the rotating dot while fixating on the center dot. After button-press or tone occurrence (depending on condition), the dot completed its current revolution and then completed one more revolution in order to avoid biasing participants’ reports. Then, participants indicated the dot’s location at the time of the relevant event (depending on the condition) by bringing the mouse cursor to the appropriate place on the clock and clicking. Although it was not used for reporting, the clock was still present on No-Report trials to keep the visual experience as similar as possible across conditions.

Our main object of investigation was to establish whether antecedent pupil dilations reflect the onset of conscious intention prior to the onset of voluntary action. We therefore had several important considerations: (1) the dilations should *not* reflect any other cognitive processes, such as reporting demands, motor execution, or general anticipation; (2) the timing of the dilations should be associated specifically with the reported time of intention awareness, but not the reported time of movement or tone awareness. Based on these considerations, we designed the experiment with five conditions, in a blocked design to make the distinction between conditions easier for participants to appreciate.

*W Condition:* Participants were instructed to wait for the clock to make a full revolution (to establish a baseline period), and then spontaneously press the spacebar on the keyboard at a time of their choice. They were specifically instructed not to pre-plan these movements, but rather to be spontaneous. In addition, they were instructed to monitor their conscious experience, noting the time (i.e., the position of the clock) when they first became aware of an urge or intention to move (note that urges and intentions are often used to refer to distinct mental states, but here we used this language to be consistent with prior studies). Participants then reported this position on the clock at the end of the trial (see Fig. 1). This condition enabled us to assess whether the timing of APDs was associated with the reported awareness of intentions.

*M Condition:* Participants were instructed to act as in the W condition, with one difference. They were now instructed to monitor their own movements and report the time (on the clock) when they pressed the spacebar (see Fig. 1). This condition enabled us to assess whether the timing of APDs was associated with the timing of action awareness.

*No-Report Condition:* Participants were again instructed to act as in the W condition, but they were also instructed not to report anything (see Fig. 1). Nor did they receive instructions to attend to their intentions, movements, or any other events. This condition enabled us to assess whether dilations were associated with reporting demands and control for those demands.

*I Condition:* Participants were again instructed to act as in the W condition, but in this condition, they were instructed to spontaneously *imagine* pressing the spacebar at a time of their choice, rather than actually press the spacebar, without pre-planning the mental action (see Fig. 1). They were further instructed to then physically press the spacebar for about half a revolution after the initial mental action of imagining the button press. This was to indicate the end of the trial (we specifically did not require precision in their estimated timing to prevent them pre-planning their action at the time of imagination). Finally, similarly to before, participants were asked to use the clock but this time to indicate when they imagined moving. This condition enabled us to control for the effects of motor execution on pupil dilations.

*S Condition:* Participants were instructed to wait until they heard a short auditory tone (PsychoPy’s default “F” tone) and then note the clock’s position at the time of the tone, without making any overt movement (see Fig. 1). Hence, they did not act spontaneously in this condition. The onset times of the tones were drawn from the participant’s response times during the initial No-Report block. This condition enabled us to assess whether APDs were associated with general anticipation, and whether the timing of APDs was associated with stimulus awareness.

### Pupil size recording and preprocessing

Pupil area (and other gaze information) was recorded online at 500 Hz using an EyeLink-1000 system, read into Python and then preprocessed using custom scripts. Preprocessing steps largely followed ref.^74^. Pupil data, while slow changing, often contain noise that must mitigated with before filtering and resampling. Hence, the first pass was over the entire pupil time-series to find NaNs (NaN = “not-a-number”) corresponding to blinks. For each blink we removed 50 ms of data before or after it to remove edge/occlusion artifacts as the eye closed or opened. Then, we removed segments where the *gaze position* deviated from fixation by 15 times or more of the mean absolute deviation. Next, we removed segments where the *dilation speed* exceeded 3 times the mean absolute deviation from median. Then, we removed segments where *pupil size* exceeded 15 times the mean absolute deviation from median. Finally, we removed pupil data that differed from a 300 ms window median (obtained using a median filter from Scipy) by more than 25 units (arbitrary units registered by the eye-tracker). Then, we linearly interpolated over every cluster of NaNs only if they were less than or equal to 600 milliseconds long, using Numpy’s 1D interpolation implementation, because longer periods likely obscured phasic changes in pupil size. Finally, to remove fast noise present in recording, we smoothed the data with a Savitsky-Golay filter with a window size of 151 ms, polyorder of 3, and extending the data for data near the edge using the nearest values (to avoid edge artifacts). Following this preprocessing, pupil data was epoched and exported for further analysis in Python.

### Statistical Analysis of Pupil Size

Trials that contained NaNs in the pupil signal after preprocessing were not considered for further analysis. After removing such trials, we had an average of 65.24 trials remaining per participant (STD: 43.71; range: 3-134). Because the number of trials remaining varied broadly across participants, we opted to use mixed-effects models (all implemented using the Pymer4 python package), which take into account single-trial information and are therefore less susceptible to adverse effects from small sample sizes than averaging signal traces within participants for each condition, and then constructing grand-averages. Furthermore, completing the same analyses as in the main manuscript on only participants with more than 40 remaining trials did not change the results discussed in the present study.

We baselined pupil size by subtracting the average pupil size on each trial in the period [-2,-1.5] s relative to movement onset. From there, we regressed the pupil size at each time point *t* on condition, with a random intercept for each participant:

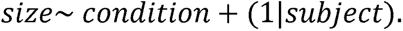

From the fitted models we obtained the estimated mean and 95% confidence intervals for each time point & condition, which are plotted in Figure 2. For comparison of pupil slope, regressed pupil size on time (−1.5 s to 0 s relative to movement, imagined movement, or tone onset) and condition (including an interaction term) and included a random intercept for each participant:

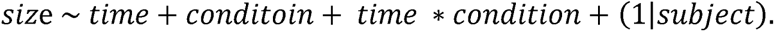

From this fitted model, we extracted the estimated interaction between time & condition, which reflects the estimated pupil slope for each condition. We then used the post-hoc tests included in the Pymer4 Package to test for significant differences between conditions.

### Analysis of Pupil Dilations and Subjective Reports

We next analyzed how pupil dilations covaried with subjective reports of times in the different conditions (using the clock). For each participant, we conducted a median-split of W, M, and S times and constructed an indicator variable (*I*_early_) for each trial that indicated whether that trial’s report was in the lower 50% (*I*_early_ = 1) or upper 50% (*I*_early_ = 0) of subjective reports (this analysis was conducted separately for the W-Time, M-Time, and S-Time conditions). No-Report and I-Time conditions were omitted from this analysis due to lack of a report or ground-truth timing for the event of interest, respectively. From there, we regressed pupil size at each timepoint t on this indicator variable with a random intercept of participant:

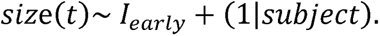

We then used post-hoc tests at each time point to determine whether the pupil size was significantly different for early vs. late reported W, M, or S time trials (cutoff □ = 0.05). Splitting based on reported W-time resulted in cluster (N = 27) of consecutive timepoints, where pupil size was significantly different across early or late W reports. Splitting based on reported M or S time did not result in any time points with significant differences (Fig. 4C).

To establish whether the size of the cluster of differences in pupil signals was significantly above chance in the W condition, we repeated the above analysis on bootstrapped data. For each participant, we shuffled whether trials were labeled as early or late, thereby retaining the participant-specific structure but destroying the statistical relation between trial identity and report. We then applied the same analysis as above and found the largest consecutive number of time points that were significantly different on each shuffle. Relative to the distribution of significant ‘clusters’ in bootstrapped data, we found that the cluster based on splitting the real data was in the top 96.4^th^ percentile of cluster sizes, suggesting that pupil dilations significantly depend on reported times of intention awareness (p = 0.036—note that cluster sizes are strictly positive, so this is analogous to a one-sided test). We did not repeat this analysis for M or S conditions because there were no significantly different clusters in those conditions. We further tested this under stricter exclusion criteria (excluding participants with fewer than 10 W-time trials) and found similar results, with a significant cluster size of 34 consecutive points, which reached the 99.07^th^ percentile across 100 bootstraps.

### Breakpoint Analysis

To investigate the timing of dilation onset, we also conducted a breakpoint analysis^75^, which seeks the onset of a change in slope. We fit multiple piecewise linear and non-linear functions to baselined pupil size waveforms between [-2 s, 0.2 s] (so that we can test breakpoints as early as-1.5 seconds) relative to movement onset in No-Report, W-Time, and M-Time trials (i.e. trials with spontaneous movements). We further fit these models only on participants who had at least 40 trials remaining after preprocessing (see above) to avoid issues of small trial numbers biasing the model fit calculations (however, fitting on all participants did not meaningfully change these results). We fit the breakpoints up to 150 ms following movement onset because after that point the dilations crest, and we are primarily interested in dilations before movement. We assumed the waveforms would be flat and then dilation would begin, so we fit the following models:

Linear:

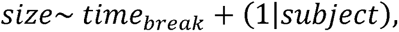

Quadratic:

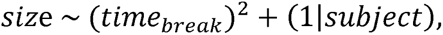

And exponential:

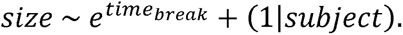

Where:

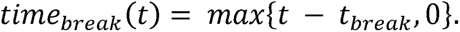

Hence, t_break_ is the time of a breakpoint (i.e., dilation onset). And time_break_ describes a piecewise linear variable that is zero before t_break_, and then increases linearly following t_break_. By varying the value of t_break_ and comparing the Akaike Information Criterion (AIC) of the fitted models, we determined the time of dilation onset that gives the best fit to the underlying data. We fit values of t_break_ between-1.5 and 0.15 s relative to movement onset (in increments of 0.05 s).

We also fit models where the slope before the breakpoint was linear rather than constant, to account for potential earlier changes in pupil size (bottom part of Fig. S3A). Those models suggested slightly later dilation onsets (between-0.8 and-0.5 seconds; Fig. S3A). Further, their AICs were lower than those of the constant fit before the breakpoint. But they visibly did not seem to capture dilation onset (Fig. S3B). Fitting models where dilation slope varied with condition brought about similar results.

### Decoding Analysis

We further wanted to investigate whether antecedent pupil dilations could be distinguished from baseline periods or time-matched periods before a non-movement event. To this end, we compared the slope of the pupil waveform at various time points before movement onset to either the distribution of slopes during the baseline period ([-2.0,-1.5 s], Fig. 4B) or to the distribution of slopes obtained from the corresponding time before tone onset in the S-Time period, where no action was generated (Fig. 4C). We used a sliding window approach (window sizes 300 ms, step size 50 ms; though the results were qualitatively similar for 100 and 500 ms windows, Fig. S3D-E). Note that the times reported in Figs. 4B and S3D-E refer to the windows’ leading edges. Further, we used the slope of the pupil waveform rather than the actual values because that avoids confounds due to baselining or consistent differences in tonic pupil size across conditions. The slopes were fit to the pupil waveform using Scipy’s linear regression algorithm. We then used linear discriminant analysis (LDA) implemented via scikit-learn to classify between (1) pupil slopes at different time points in conditions with spontaneous movements (No-Report, W-Time, and M-Time conditions) against a distribution of pupil size obtained from a baseline period long before movement (starting at-2.0 s relative to movement using the window size later used for decoding; Fig. S3C) and (2) pupil slopes during conditions with spontaneous movements (No-Report, W-Time, and M-Time conditions) and a window with the same leading edge without movements (S-Time condition). Note that these data were thus a 1-dimensional input for these decoding algorithms, which are usually used for multidimensional data, but we employed them to compare against studies that try to predict movement from other signals e.g. EEG. For each fitted model, we calculated the average AUC at each time point across 3 cross-validations for each participant. We used AUC instead of accuracy because decoding vs. the S condition involved an unbalanced dataset (more spontaneous movement trials than S-Time trials). And we used the relatively low 3-fold cross validation due to the low trial numbers (AUC can only be calculated when a randomly chosen validation set has at least one of each type of trial). We next performed the same decoding analysis on data pooled across participants, randomly shuffling the data 1000 times (pooled to make sure that all random shuffles would result in enough samples for AUC calculation for all individual participants) to obtain a chance distribution for AUC. Note that such machine-learning techniques are usually used for high-dimensional input spaces, whereas here we only use pupil slope (hence a 1D input space). We do this for two reasons: first, to assess *predictiveness* of pupil dilation as opposed to just differences prior to movement onset; second, to compare to other analyses trying to predict upcoming movement from neuroimaging data, which is higher-dimensional than pupil size.

## Data and Code Availability

Data and analysis code will be made available upon publication of the article.

## Author Contributions

**Jake Gavenas**: conceptualization, methodology, software, formal analysis, investigation, visualization, writing – original draft; **Aaron Schurger**: methodology, resources, writing – review & editing, supervision; **Uri Maoz**: conceptualization, methodology, validation, resources, writing – review & editing, supervision, funding acquisition.

## Supplementary Figures

**Supplementary Figure 1.**
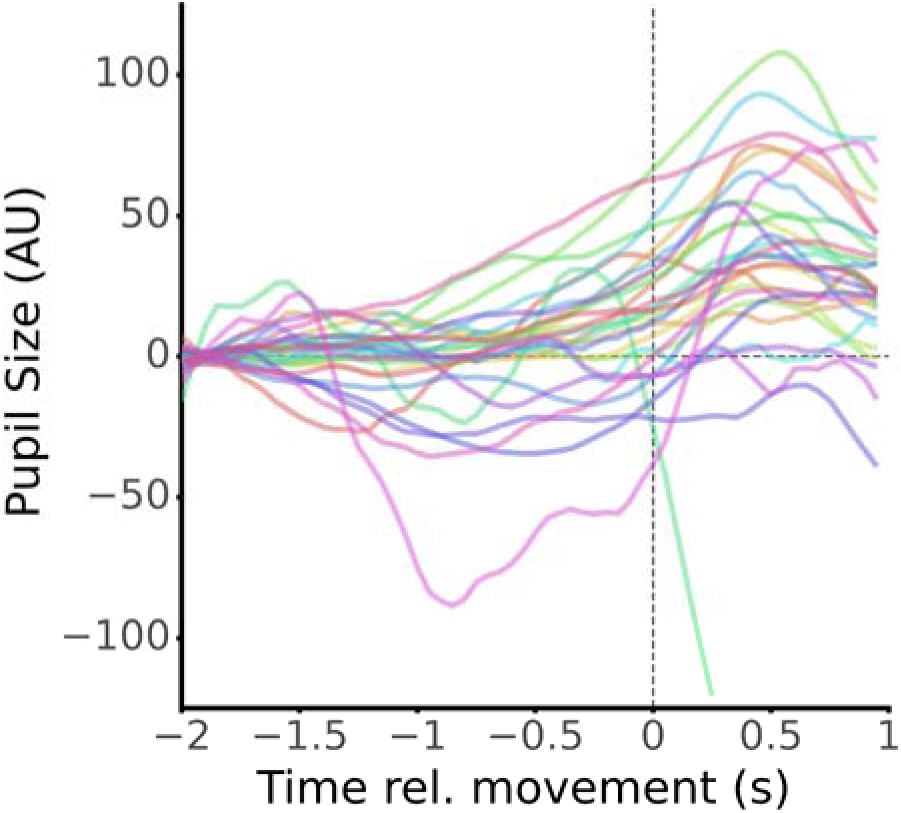
Antecedent pupil dilations from individual participants. Most participants showed a gradual dilation in the time leading up to movement, with a few exceptions.

**Supplementary Figure 2.**
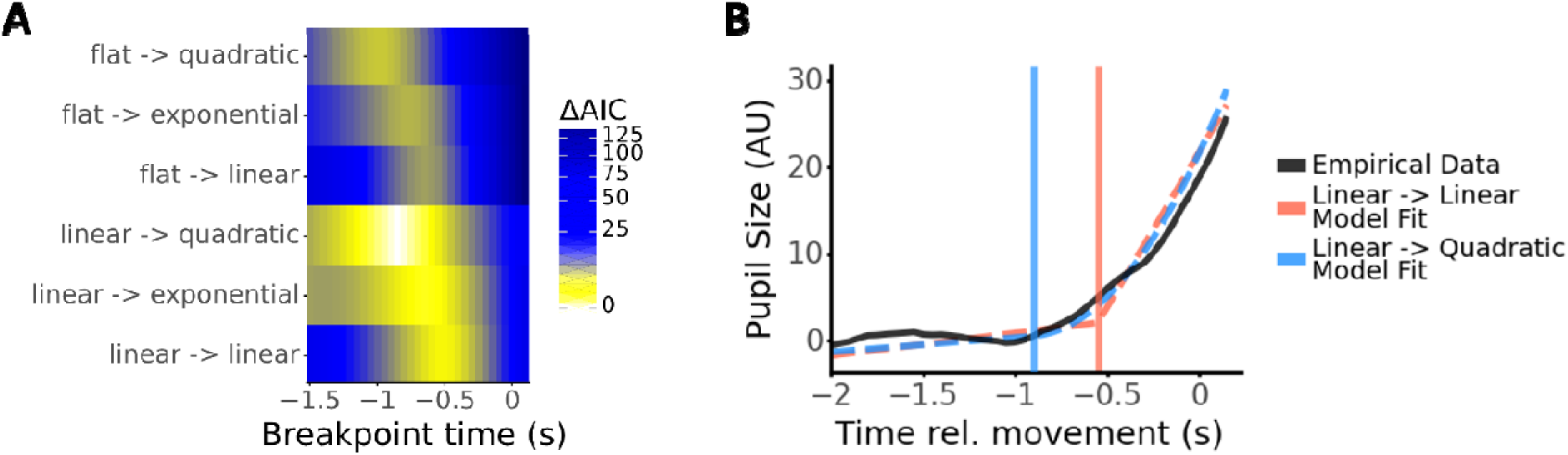
Supplementary results from breakpoint & decoding analyses. **A:** Breakpoint analysis for piecewise functions that had a linear component and then a linear, quadratic, or exponential component (the functions before and after the breakpoint are separated by the “->” symbol). “Flat” (in the top half) refers to models where the initial component was constant (i.e., linear that was forced to have zero slope), and “linear” (in the bottom half) refers to models where the initial component linear could have slope different from zero. Plotted is a heatmap of AICs relative to the best-performing model as in Fig. 3A, except for all model types (with a flat/constant and linear pre-breakpoint fit; note that the “flat-> linear” model here corresponds to the “linear” model referred to in the main text. **B:** Model fits using the breakpoint that resulted in lowest AIC for the linear-> linear and linear-> quadratic models (linear-> exponential was omitted as it largely overlapped with the quadratic model). Solid line is grand-average pupil size (averaged across trials and then across participants), dashed lines are model fits, and vertical solid lines are the breakpoints used for the corresponding models (as in Fig. 3B).

**Supplementary Figure 3.**
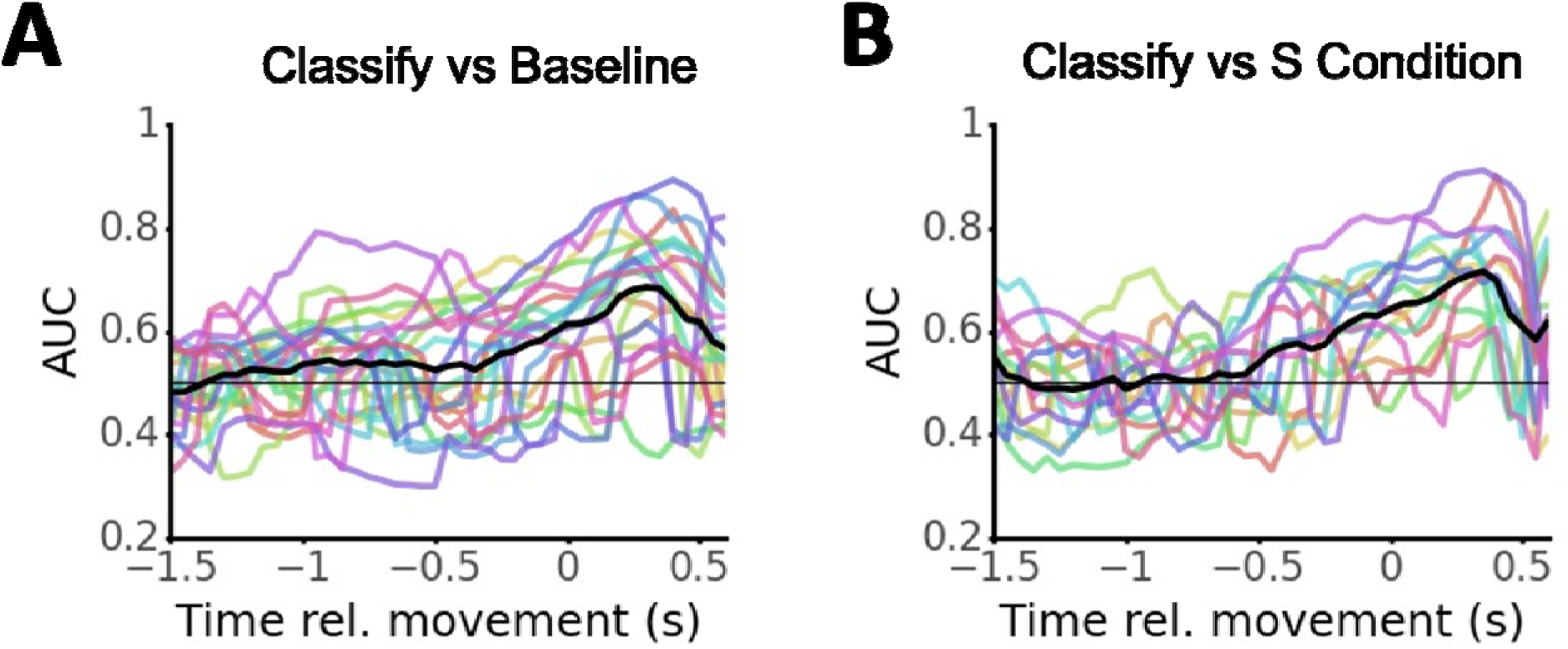
Decoding performance for individual 995 participants. Colored lines are individual traces of decoding test-set AUC over time (0.3 s sliding window; average of 3-fold cross validation). Black line is average over all participants. **A.** Decoding versus baseline. **B.** Decoding versus S condition.

**Supplementary Figure 4.**
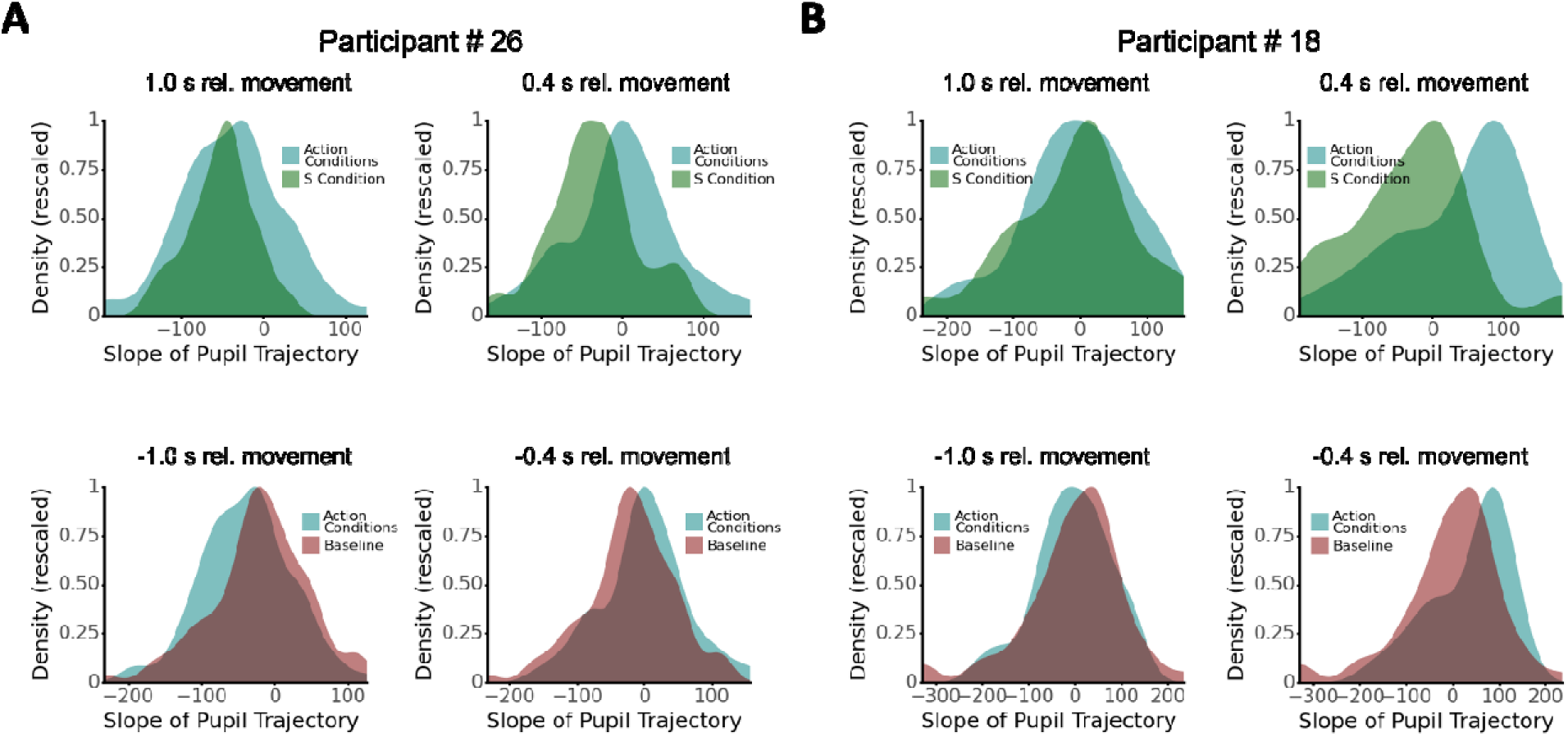
Shifts in distribution of pupil slope underlie decoder performance. Density plots of pupil slopes (0.3 s window; densities calculated using Plotnine’s default method using geom_density; rescaled so peak value is 1 for better comparison) for two typical participants for decoding action condition (No-Report, W, M) from S condition (top row) or baseline (bottom row). 1 second from movement the distributions are highly overlapping (left columns in panels A and B), but the distributions for slope during action conditions shifts rightward (more positive slope indicating pupil dilation) closer to time of movement.

**Supplementary Figure 5.**
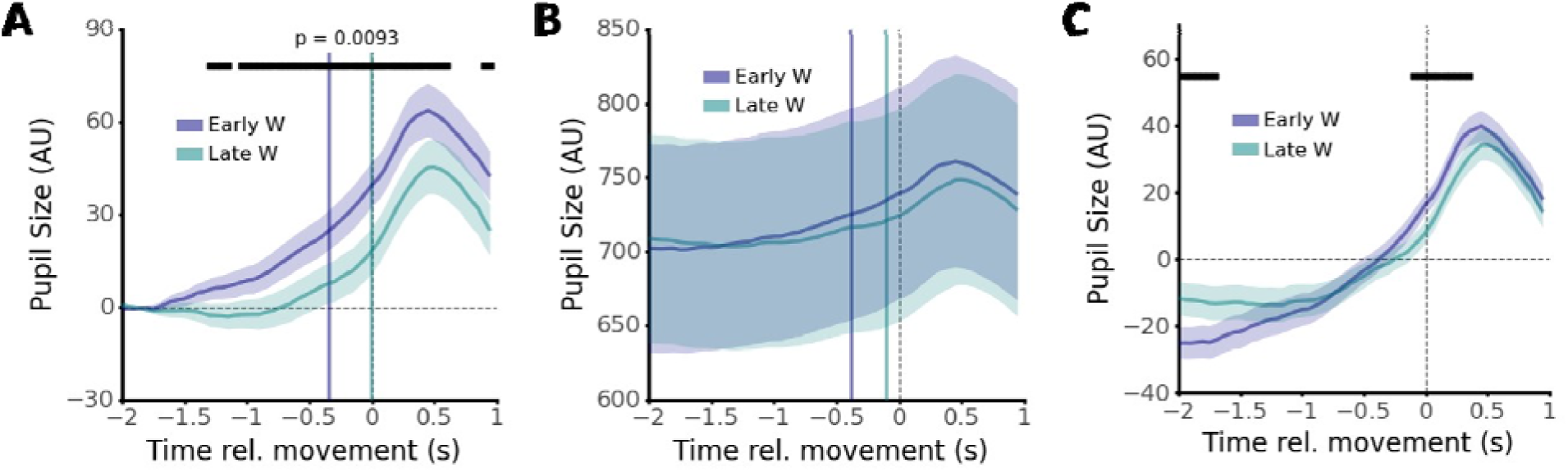
Further analysis of relation between early & late W reports and pupil waveform. **A.** Replicating Fig. 4A with stricter exclusion criteria (>10 trials of W-time condition per participant), we found that earlier reported W-times were still significantly associated with earlier dilations (p = 0.009, obtained from non-parametric cluster permutation test, actual cluster size = 34 consecutive timepoints, 99^th^ percentile of shuffled data (N=100 bootstraps). **B.** Replicating Fig. 4A on non-baselined pupil data. Due to the lack of baseline correction, confidence intervals are much wider than baseline corrected versions, leading to no significant difference between conditions. However, dilations are visibly present for both trials, and occur earlier for early W trials. Notably, dilations on early W trials also reach a visually larger peak dilation compared to late W trials. **C.** Replicating Fig. 4A on pupil data that was demeaned by subtracting the whole-trial average from each time-point. LME analysis suggests significant differences between pupil size early in the trial (−2 to around-1.6 s relative to movement) and around the time of movement, which is consistent with dilations on early W trials being earlier and stronger.

